# Ubiquitin Chloromethylketone Probe Enables Activity-based, Selective Protein Profiling of E2 Ubiquitin Conjugating Enzymes

**DOI:** 10.64898/2026.06.16.732757

**Authors:** Saibal Chanda, Alan Pham, Yugendar R. Alugubelli, Nathan Axtell, SreeNidhi Karnati, Yashwanth Aravapalli, James X. Mao, Wenyue Cao, Wenshe Ray Liu

## Abstract

Ubiquitin (Ub) conjugating enzymes (E2s) are central to Ub signaling, yet their systematic activity-based profiling remains challenging due to the weak nucleophilicity and elevated pKa of their catalytic cysteines. Existing Ub probes primarily target deubiquitinases (DUBs) and the only reported E2-targeting probe requires E1-dependent activation to capture limited E2s. To profile E2s broadly, here Ub chloromethylketone (UbCMK) is reported as a standalone activity-based probe. Density functional theory calculations identified CMK as a highly electrophilic warhead with a low activation barrier for reaction with weakly nucleophilic thiolates. UbCMK was synthesized via activated cysteine-based protein ligation and irreversibly labeled multiple E2s and cysteine DUBs. Activity-based protein profiling and quantitative proteomics in HEK293T cell lysates revealed broad enrichment of E2 enzymes, including many previously inaccessible to other probes. UbCMK furthermore enables activity-dependent quantification of endogenous E2 mobilization across oxidative, proteotoxic, inflammatory, metabolic, lipid oxidative, and genotoxic stress conditions. In addition, UbCMK engages both E1s and DUBs as well, indicating its broad utility as a probe. Collectively, these results establish UbCMK as a powerful chemical tool that expands activity-based protein profiling coverage across the Ub-proteasome system and enables functional interrogation of E2 enzymes under physiological and pathological conditions.

## INTRODUCTION

Ubiquitin (Ub) is an evolutionarily conserved 76-amino acid protein present in nearly all eukaryotes.^1^ Its covalent attachment to other proteins, commonly known as protein ubiquitination, regulate their function, localization, and/or homeostasis.^2,3^ The most notable function of protein ubiquitination is to mark proteins for proteasomal degradation.^4^ The Ub-proteasome system (UPS) thus governs protein homeostasis and the regulation of many fundamental cellular pathways,^5^ and its dysregulation has been linked to multiple diseases.^6^ The development of PRoteolysis-TArgeting Chimeras (PROTACs) that uses the UPS to drive degradation of therapeutic target proteins has also revolutionized the drug discovery field, leading to many PROTACs on clinical trials.^7^ Protein ubiquitination is catalyzed by an enzymatic cascade comprising three enzymes including Ub activating enzyme E1, Ub conjugating enzyme E2, and Ub ligase E3 (Figure 1A), which act concertedly to activate and attach the Ub *C*-terminal carboxylic acid to a lysine side chain amine on a substrate protein by forming an isopeptide bond. This modification is reversed by deubiquitinases (DUBs), thereby preserving the dynamic balance of Ub signaling pathways. To characterize enzymes functioning in Ub signaling pathways, many Ub-based chemical probes have been developed (Figure 1B). The earliest Ub probe, Ub aldehyde, inactivates DUBs with a catalytic cysteine by forming a thiohemiacetal adduct.^8^ Ub nitrile, developed subsequently, inhibits a 26S proteasomal isopeptidase through thioimidate formation.^9^ Since both probes lead to reversible covalent adducts with DUBs, other probes incorporating Michael acceptors or alkyl halides to form stable adducts with DUBs were later developed.^10^ Interestingly, although a typical Ub Micheal acceptor probe doesn’t efficiently engage E2 enzymes that contain an active site cysteine as well, a Ub probe with a *C*-terminal dehydroalanine (UbDHA) was reported to react with E2.^11^ Its free *C*-terminal carboxylic acid allows its activation by E1 via an ATP hydrolysis-driven process to form an E1 complex that binds more potently to E2 and therefore brings its dehydroalanine moiety close to the E2 cysteine for a proximity-driven reaction. Many other Ub probes that allow activity-based protein profiling of enzymes in the UPS have also been developed. A notable one is Ub propargylamine (UbPA).^12^ Due to concerns that certain DUB family enzymes such as OTU DUBs possess narrow catalytic clefts with stringent geometry for strict substrate preferences and thus display weak reactivity toward some conventional probes,^13^ we developed Ub azapeptide ester probes featuring an azaglycine ester at the Ub *C*-terminus to closely mimic native substrates for UPS enzymes.^14, 15^ These new probes provide better activities toward certain DUBs than UbPA but are still lack of efficiency to be generally applied as E2 probes.

**Figure 1.**
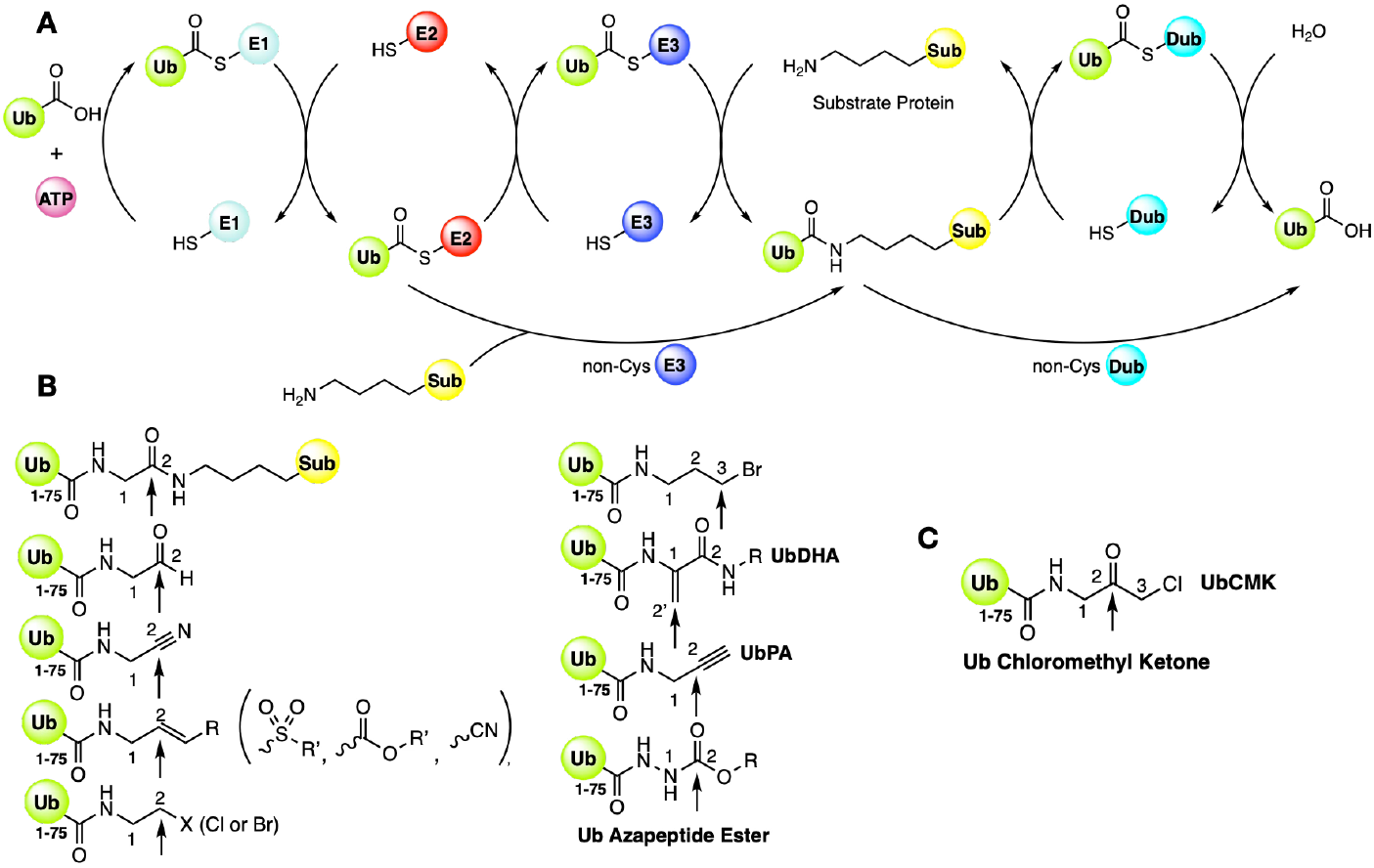
The ubiquitin (Ub) signaling pathway and Ub probes. (**A**) The E1-E2-E3 cascade mediates the covalent attachment of Ub to a substrate protein (Sub), the process begins with an ATP dependent activation of Ub by an E1activating enzyme, formingE1-Ub thioester. The activated Ub is then transferred to E2 conjugating enzyme via transthiolation and finally ligated to the substrate via E3 ligase. Deubiquitinases (DUBs) catalyze the reverse process. Both E3 ligases and DUBs can be either cysteine-dependent or cysteine-independent. (**B**) Existing Ub probes shown alongside a ubiquitinated protein; Gly76 Cα (1) and carbonyl carbon (2) are indicated, and arrows mark the scissile bond in Ub-Sub as well as the electrophilic positions in each probe that can form covalent adducts with catalytic cysteines in the Ub signaling pathway enzymes. (**C**) Proposed Ub chloromethylketone (UbCMK) probe for E2 conjugating enzymes.

So far, only UbDHA has been used as a probe for activity-based protein profiling (ABPP) of E2 enzymes.^11^ Since UbDHA needs to be activated by E1 in order to engage E2, how exactly the dehydroalanine residue at the Ub *C*-terminus affects the activation process catalyzed by different E1 enzymes is unknown. The requirement of the formation of an activated E1-UbDHA complex for engaging E2 potentially complicates the ABPP analysis as well since different cells and tissues express different levels of E1 enzymes. However, studying E2 has become increasingly significant. In cancer, aberrant E2 activity can contribute to progression by sustaining proliferation, genomic stability, immune invasion, metastasis, and treatment resistance.^16^ For example, UBE2C and UBE2S are frequently associated with mitotic control and poor prognosis,^17^ UBE2T has been linked to Franconi anemia pathway regulation, epithelial-mesenchymal transition, PI3K/AKT and Wnt/β-catenin signaling, and multiple tumor types,^18^ and UBE2N/UBC13 supports K63-linked Ub signaling involved in NF-κB activation, DNA damage responses, and more recently, acute myeloid leukemia proteostasis.^19^ Therefore, a generally applicable ABPP probe for E2s will potentiate accelerated systematic understanding of E2 functions in cellular and developmental biology and therapeutic candidates targeting them. In this work, we wish to report such a probe.

## RESULTS and DISCUSSIONS

### UbCMK as a Potential ABPP Probe for E2 Enzymes Based on DFT Calculation

Many Ub-based ABPP probes fail to efficiently capture E2 enzymes due to several key factors. First, the pKa of E2 active-site cysteines is elevated by approximately 2 pH units above that of a free cysteine (∼10.5 vs. ∼8.5 for free Cys).^20^ This elevated pKa could be a regulatory mechanism which prevents the exposed E2 active-site cysteine from reacting promiscuously with electrophiles in the cell. Secondly, most E2s bind weakly to Ub.^21^ Thirdly, there is fundamental mechanistic mismatches among cysteine enzymes in the UPS. As shown in Figure S2, DUBs need to catalyze the hydrolysis of a strong isopeptide bond between Ub and a substrate protein and therefore requires a strongly nucleophilic active site cysteine activated by a histidine as its catalytic dyad partner. E1 catalyzes two consecutive reactions with the first as the formation of Ub-AMP followed by the transfer of Ub from Ub-AMP to its active site cysteine. Ub-AMP has an activated Ub ester bond for reaction with the E1 cysteine. A strong nucleophilic cysteine such as in a catalytic dyad is not necessary for this reaction but E1 enzymes such as Uba1 still have a nearby arginine residue to deprotonate the active site cysteine for a reaction exchange with AMP in Ub-AMP. Unlikely a DUB and E1, E2 receives Ub from E1-Ub via a thiol exchange reaction that requires a low or no activated E2 cysteine. An E2 active site cysteine, e.g. in UBE2F, is typically solvent exposed for maintaining its low nucleophilic feature.^22^

To develop an E2-targeting probe, we thought to replicate the thiol exchange reaction that requires low activation energy. Therefore, we conducted DFT calculation of chemical groups including azapeptide ester, alkyne in propynyl acetamide, vinylsulfone, and chloromethylketone (CMK), shown in Figure 2A, on their nucleophilicity and activation energy to react with a thiolate. CMK was included in this calculation because it was previously used in covalent inhibitors for both cysteine and serine proteases. A well accepted mechanism for a CMK probe that was confirmed by NMR for reacting with either cysteine or serine in an enzyme is through a tetrahedral thiohemiacetal or hemiacetal intermediate (Figure S1).^23^ The thiohemiacetal intermediate then undergoes a 3-center reaction to remove chloride for generating a sulfonium ion that rearranges to form the finally stable thiomethylketone adduct. Our calculation results show that CMK displays the highest electrophilicity among all four chemical groups at its carbonyl carbon for engaging a thiolate (Figure 2B). The calculated Gibbs free energy of activation for the four functionalities as shown in Figure 2C clearly indicated that CMK has significantly lower activation energy than the other three chemical groups for reacting with a thiolate, making it more likely to engage an E2 active site cysteine. Encouraged by these calculation results, we proceeded to synthesize a UbCMK probe with the Ub *C*-terminal glycine substituted by CMK.

**Figure 2.**
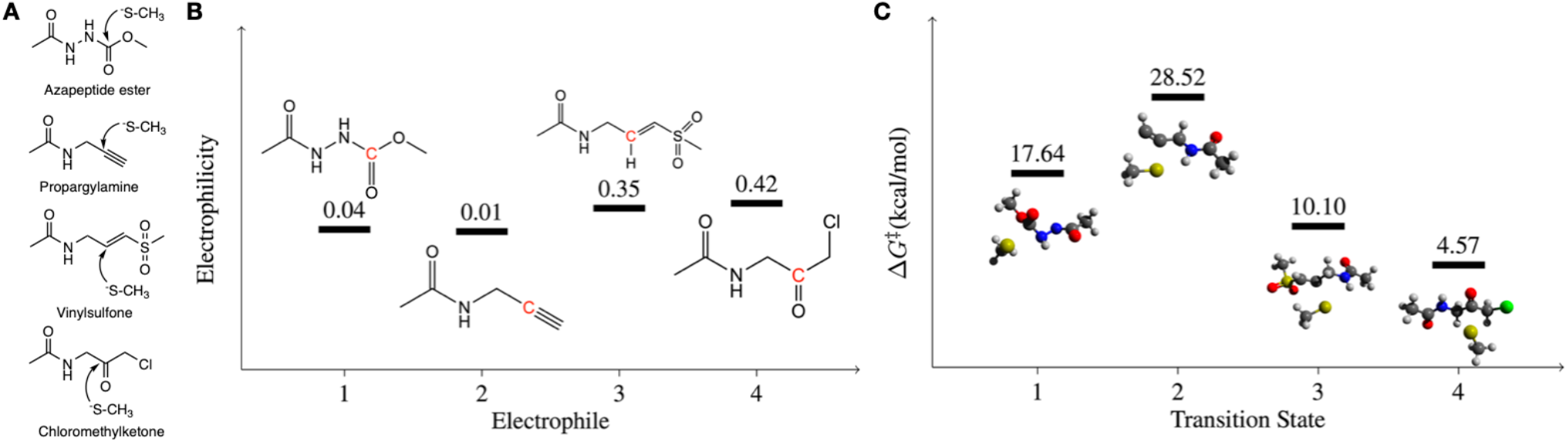
DFT-guided identification of CMK as a potential electrophile for E2 active-site cysteines. (**A**) Schematic diagrams showing cysteine thiolate reacting with different chemical groups (azapeptide ester, propargylamine, vinyl sulfone and CMK) used for DFT calculation. (**B**) Electrophilicities thio-reacting C atoms from various electrophiles, calculated by Fukui function: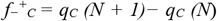, where *qC (N)* is the NPA charge of C atom and N is the number of electrons in the neutral system. Calculations were carried out at the M06-2X/6-311+G(d,p) level of theory, with the PCM model employed to account for solvation effects. (**C**) Calculated Gibbs free energies of activation for the addition reactions of CH3S^-^ and various electrophiles. Calculations were carried out at the M06-2X/6-311+G(d,p) level of theory, with the PCM model employed to account for solvation effects.

### Synthesis of FLAGUbPA and FLAGUbCMK

Expressed protein ligation is a widely used technique for the synthesis of Ub probes.^24^ An alternative activated cysteine-based protein ligation (ACPL) technique that utilizes a nitrile donor reagent such as 2-nitro-5-thiocyanatobenzoic acid (NTCB) to activate a cysteine residue within the protein for an exchanging reaction with a small-molecule amine has also been used to synthesize Ub probes (Figure 3A).^14, 25^ Due to its simplicity, we used ACPL to generate Ub probes in the current study. FLAG-Ub_1-75_-Cys-6×His with a *N*-terminal FLAG tag (DYKDDDDK), a G76C mutation, and a *C*-terminal 6×His tag was expressed and purified from *E. coli* (Figure S1). We followed a previously published procedure to synthesize FLAGUbPA (FLAG-Ub_1_-_75_-propargylamine). To synthesize FLAGUbCMK (FLAG-Ub_1_-_75_-chloromethylketone), we followed a two-step chemical reaction strategy shown in Figure 3B. FLAG-Ub_1-75_-Cys-6×His was first reacted using the ACPL method with small-molecule amines such 3-chloro-2,2-diethoxypropan-1-amine to yield a corresponding Ub-amine conjugate that subsequently underwent acid-catalyzed acetal deprotection to afford FLAGUbCMK. This two-step synthesis scheme is required because 1-amino-3-chloroketone reacts to itself at a high concentration and cannot be applied to the ACPL reaction. For a comparison purpose, we also generated FLAGUbFMK (FLAG-Ub_1-75_-fluoromethylketone) that has a less reactive fluoride leaving group. Throughout this study, FLAGUbPA served as a control molecule due to its widespread use as an activity-based probe for DUBs. All three Ub probes were purified to homogeneity using fast protein liquid chromatography (FPLC) (Figure S3). The purified products were subsequently characterized by electrospray ionization mass spectrometry (ESI-MS) to confirm their identities and purity. For quality control, the precursor protein FLAG-Ub_1_-_75_-Cys-6×His was also analyzed under identical conditions. Representative proton-charged and deconvoluted ESI-MS spectra for all constructs are provided in Figures S4-S7, with the deconvoluted spectra of the final probes summarized in Figure 3C, along with their experimentally determined molecular weights. Across all probes, the determined molecular masses were in excellent agreement with their theoretical values, exhibiting a maximum deviation of 0.3 Da (Table S1). These results collectively verify the successful synthesis and structural integrity of three Ub probes. To assess whether FLAGUbCMK and FLAGUbFMK retain a native-like secondary structure, circular dichroism (CD) spectroscopy was performed. The CD spectrum of FLAGUbCMK and FLAGUbFMK closely resembled that of FLAG-Ub_1-75_-Cys-6×His (Figure 3D), indicating that the overall protein fold is preserved.

**Figure 3.**
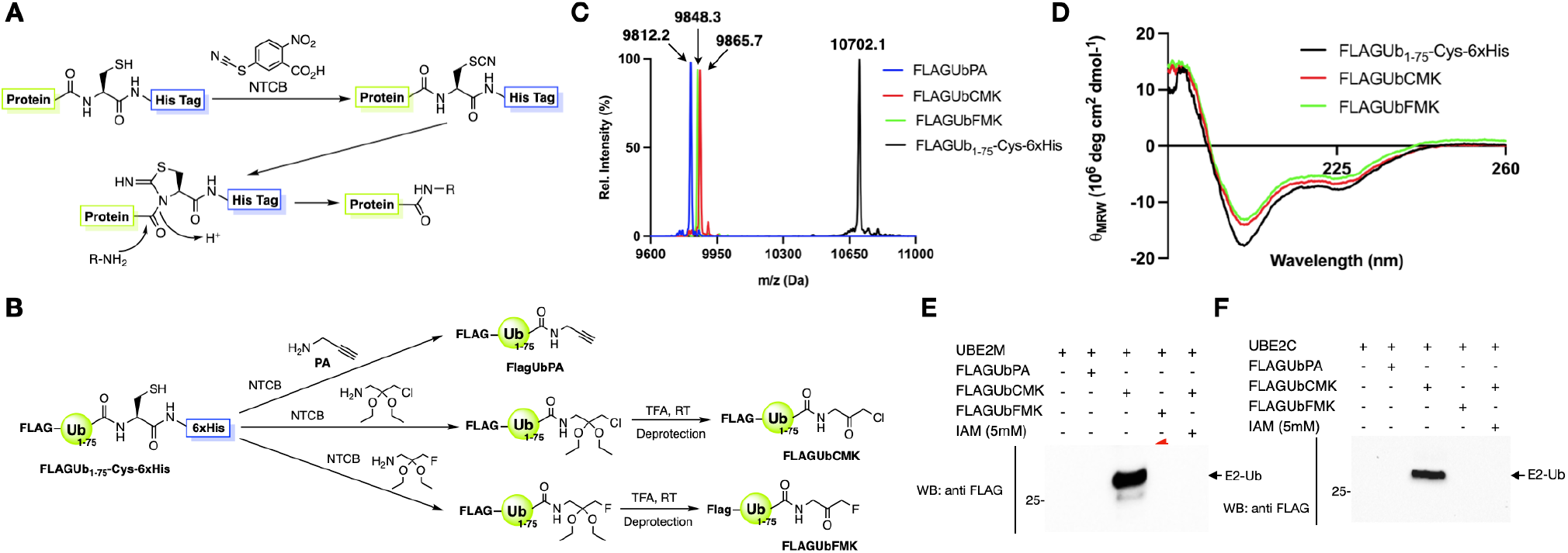
Synthesis of FLAGUbPA, FLAGUbCMK, and FLAGUbFMK using ACPL. (**A**) Schematic diagram of the ACPL reaction mechanism used to functionalize the *C*-terminal cysteine of FLAGUb_1-75_-Cys-6×His. The process involves activation of the cysteine by NTCB to form a 1-acyl-2-iminothiazolidine intermediate, followed by nucleophilic acyl substitution with small molecule amines to afford the desired Ub conjugates. (**B**) The synthetic routes for generating FLAGUbCMK, FLAGUbFMK from FLAGUb_1-75_-Cys-6×His. Reactions include (i) coupling with 3-chloro-2,2-diethoxypropan-1-amine and 3-fluoro-2,2-diethoxypropan-1-amine, respectively, followed by acid-catalyzed deprotection. FLAGUbPA was synthesized as a control probe for the study. (**C**) Deconvoluted ESI-MS spectra of three final Ub probes, with observed molecular weights closely matching theoretical values (Δ ≤ 0.3 Da), verifying the identity and purity of the synthesized probes. (**D**) CD spectra of FLAGUbCMK, FLAGUbFMK, and FLAGUb_1-75_-Cys-6×His which shows that both halomethyl ketone probes retains a secondary structure consistent with FLAGUb_1-75_-Cys-6×His, indicating proper folding after the ACPL and deprotection reactions. (**E**-**F**) Covalent complex formation of UBE2M and UBE2C, respectively, with FLAGUbCMK in contrast to unreactive FLAGUbPA and FLAGUbFMK towards the two tested enzymes. The formed E2-Ub complexes are indicated. The complex formation was blocked by iodoacetamide confirming the engagement of catalytic cysteines of the enzymes.

### Validation of FLAGUbCMK as a E2 Covalent Probe

With the successful synthesis of FLAGUbCMK, we next evaluated its ability to covalently label an E2 enzyme. UBE2M was selected as a model enzyme for this validation and both FLAGUbPA and FLAGUbFMK were included in this validation test for comparisons. Recombinant UBE2M was incubated with one of the three probes under identical conditions (60 min at 37 °C) and then the reaction mixtures were analyzed by SDS-PAGE and western blotting using an anti-FLAG antibody. As shown in Figure 3E, UBE2M itself (lacking a FLAG tag) exhibited no detectable signal, whereas incubation with FLAGUbCMK produced a higher molecular weight adduct (∼32 kDa), consistent with the UBE2M-Ub complex formation. Interestingly, both FLAGUbPA and FLAGUbFMK failed to produce a detectable UBE2M-Ub complex. To confirm equal protein loading, reactions were performed in a standard reaction buffer supplemented with bovine serum albumin (BSA) as an internal control, which was verified using an anti-BSA antibody (Figure S9). To confirm the labeling arises from the enzyme’s catalytic cysteine, we treated the enzyme with 5mM iodoacetamide followed by incubation with FLAGUbCMK. As shown in Figure 3E, pre-alkylating catalytic cysteine completely abolished the complex formation supporting the involvement of enzyme’s catalytic cysteine. The fact that UBE2M reacts readily with FLAGUbCMK but not FLAGUbFMK can be explained by the slowing leaving group nature of the fluoride atom. A similar characterization was also conducted with another E2 enzyme, UBE2C. Similar observation was made. UBE2C reacted only with FLAGUbCMK in the reaction time frame and the UBE2C-Ub complex formation is dependent on its catalytic cysteine (Figure 3F).

### Validation of FLAGUbCMK as a general Covalent Probe for DUBs

To assess how the CMK warhead behaves toward canonical cysteine protease-like active sites, we characterized reactions of FLAGUbCMK with a panel of cysteine DUBs and compared with FLAGUbPA and FLAGUbFMK. 10 DUBs were tested, Including two from the UCH family (UCHL1, UCHL3), six from the USP family (USP2, USP5, USP7, USP9X, USP10, USP14), and one each from the Josephin family (Ataxin-3) and OTU family (OTUD6B). Reactions of all DUBs with three probes were conducted and analyzed in the same manner as for UBE2M. Molecular weights of each DUB and its covalent complexes with three probes are listed in Table S2A. As shown in Figure S10A-D, all three Ub probes gave robust labeling across multiple DUBs, producing distinctly higher molecular weight bands consistent with covalent DUB-Ub adducts. For all tested DUBs, FLAGUbCMK exhibited reactivity comparable to or greater than FLAGUbPA. FLAGUbFMK displayed a weaker and more restricted labeling profile compared to the other two probes. Comparatively, FLAGUbCMK is the most broadly and strongly reactive toward this DUB panel. Results so far demonstrate that FLAGUbCMK can serve as a general covalent probe for both E2 and DUB enzymes.

The combined E2 and DUB reaction data provide a clear framework for evaluating warhead performance and inform the choice of probes for subsequent studies. FLAGUbCMK consistently performs at or above the level of FLAGUbPA and FLAGUbFMK across the DUB panel and, crucially, is the only probe that efficiently labels UBE2M and UBE2C. In light of this superior performance and broader applicability, we elected FLAGUbCMK as a broadly applied ABPP probe for cysteine enzymes in the UPS and discontinued further exploration with FLAGUbFMK.

### Characterization of FLAGUbCMK Covalent Complexes with UPS Enzymes

When FLAGUbCMK encounters the active site cysteine of an E2 enzyme, a multi-step reaction mechanism is involved to generate a thiomethylketone covalent complex. A final irreversible thioether adduct is formed. To characterize the adduct formation and its influence by different conditions, we selected UBE2D2 as a model enzyme (Figure 4A). We characterized and assessed the stability of the thioether complex formed between UBE2D2 and FLAGUbCMK under a range of temporal, thermal, and redox conditions. First, we evaluated the stability of the complex over time. UBE2D2 and FLAGUbCMK at a 1:1 ratio was premixed for 1 h min and then tested at different conditions. Continuous incubation at 37 °C for up to 24 h did not result in detectable loss of the covalent adduct (Figure 4B), demonstrating excellent temporal stability under physiologically relevant conditions. To evaluate its thermal stability, the UBE2D2-FLAGUbCMK thioether complex was incubated at increasing temperatures (37, 45, 50, 60, 80, and 100 °C) for 60 min. As shown in Figure 4C, the complex remained fully intact up to 80 °C, with only a slight decrease in signal observed at 100 °C, indicating that the thioether linkage has excellent thermal stability. We further probed whether the covalent thioether bond was susceptible to thiol exchange in a reducing intracellular environment. A thiol competition assay was performed by incubating the complex with increasing concentrations of glutathione for 60 min. As shown in Figure 4D, the complex remained largely stable across all tested glutathione concentrations, with only a minimal decrease at 10 mM, a concentration exceeding typical cytosolic conditions. Finally, to confirm that the covalent labeling occurs specifically through the active-site cysteine of UBE2D2, the enzyme was pre-treated with iodoacetamide prior to incubation with FLAGUbCMK. Pre-alkylation completely abolished complex formation (Figure 4E), confirming that the covalent bond is formed via the active site cysteine and results in a stable thioether linkage. Collectively, these data demonstrate that FLAGUbCMK forms a robust and stable covalent complex with an E2 enzyme. The observed thermal, temporal, and redox stability highlights its suitability as a reliable ABPP probe for profiling E2 enzymes in the UPS.

**Figure 4.**
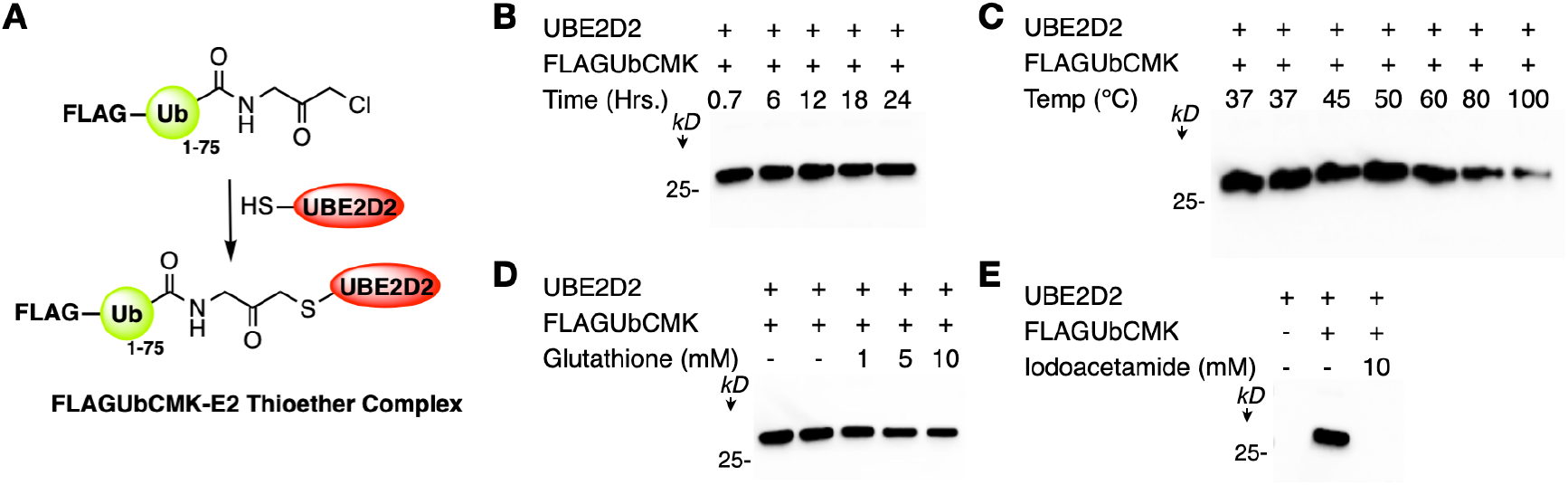
Characterization of the covalent thiomethylketone-connected UBE2D2-Ub complex. (**A**) Structure of a UBE2D2-FLAGUbCMK thiomethylketone complex. (**B**) The stability of the complex over time at 37 °C. (**C**) Stability of the complex at different temperatures for 60 min. (**D**) Stability of the complex under varying concentration of glutathione for 60 min. (**E**) The complex formation is completely abolished by pre-treating the enzyme with iodoacetamide, reflecting the involvement of enzyme’s catalytic cysteine in covalent bond formation with the probe.

### Activity-based protein profiling using FLAGUbCMK

To evaluate FLAGUbCMK in its capability to profile cysteine enzymes in the UPS, we performed activity-based labeling experiments using HEK293T cell lysates. Clarified lysate supernatants were incubated with FLAGUbPA (control) and FLAGUbCMK under identical conditions (20 h, 4 °C). To minimize batch-to-batch variability, the same lysate preparation was used for all labeling reactions. Reaction mixtures were analyzed by SDS-PAGE followed by western blotting using an anti-Flag antibody. In the absence of a probe, no anti-FLAG detected band was observed. In contrast, incubation with both probes resulted in the formation of numerous Ub-conjugated adducts that were robustly detected by anti-FLAG (Figure 5A), indicating broad labeling of cysteine enzymes. However, under a same exposure time, FLAGUbCMK-treated samples displayed significantly more labeled protein complexes and higher labeling intensity for the majority of the complexes. There were several detected bands in both samples with similar molecular weights, consistent with labeling of common enzyme populations. However, significantly more bands with enhanced labeling intensity were observed in the FLAGUbCMK-treated sample but were either weak or undetectable in the FLAGUbPA-treated sample. Many prominent bands at above 37 kDa were strongly enriched in the FLAGUbCMK-treated sample, whereas only weak labeling was detected with that for FLAGUbPA. Two intense bands present around 50 kDa and slight above were uniquely observed in the FLAGUbCMK-treated sample and were notably absent in the FLAGUbPA treated sample, indicating selective capture of distinct cysteine enzymes by the CMK warhead. At higher molecular weights, several bands in the 75-100 kDa range were prominently labeled in the FLAGUbCMK-treated sample but showed reduced or undetected intensity in the FLAGUbPA-treated sample. Collectively, these labeling results indicate that while both probes label a common subset of enzymes, FLAGUbCMK exhibits enhanced labeling efficiency and broader coverage across the UPS proteome. This increased labeling is consistent with the higher intrinsic electrophilicity of the CMK warhead, enabling more efficient covalent engagement of cysteine enzymes. As a negative control, HEK293T lysates were incubated with FLAG-Ub_1_-_75_-Cys-6×His, which lacks an electrophilic warhead. No significant labeling was observed (Figure S12), confirming that covalent capture requires an activated *C*-terminal electrophile and supporting the specificity of the labeling reactions.

**Figure 5:**
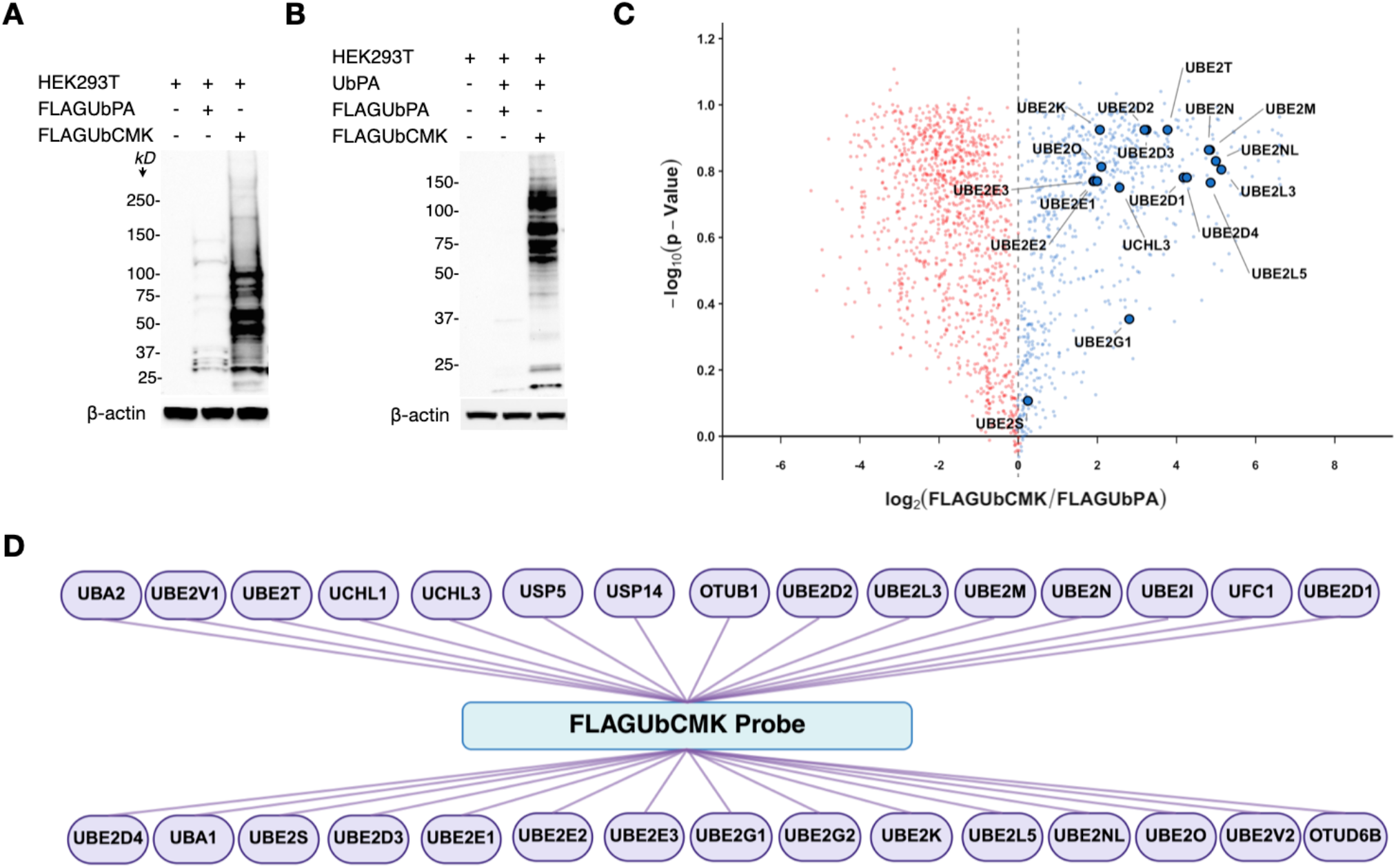
Activity-based protein profiling in HEK293T using the UbCMK probe. (**A**) Cysteine enzymes in HEK293T cell lysates were profiled by FLAGUbPA and FLAGUBCMK, respectively, via incubating lysates with each probe for 20 h, followed by SDS-PAGE and anti-Flag western blot analyses. (**B**) Cysteine enzymes from HEK293T cells were selectively profiled by FLAGUbCMK by first incubating cell lysates with UbPA for 12 h to remove UbPA-reactive enzymes, followed by a second 12 h incubation with FLAGUbPA and FLAGUbCMK, respectively, before SDS-PAGE and anti-Flag western blot analyses. (**C**) Volcano plots showing the relative enrichment of E2 enzymes in UbPA-pretreated HEK293T lysates following secondary labeling with FLAGUbCMK, plotted as log_2_(FLAGUbCMK/FLAGUbPA) versus -log_10_(p value), with significantly enriched E2 enzymes and one DUB highlighted and annotated. (**D**) Ub signaling pathway enzymes identified from FLAGUbCMK treated samples in **B.**

To further identify enzymes that may be uniquely targeted by FLAGUbCMK, we implemented a competitive blocking strategy using a non-FLAG-tagged UbPA probe to selectively inactivate UbPA-reactive enzymes prior to labeling with FLAGUbCMK. UbPA was synthesized by ACPL from Ub_1_-_75_-Cys-6×His and confirmed by ESI-MS (Figures S13 and S14). HEK293T lysates were pretreated with UbPA for 12 h at 4 °C and subsequently incubated with FLAGUbPA or FLAGUbCMK for an additional 12 h. Samples were then analyzed by SDS-PAGE and anti-FLAG western blotting (Figure 5B). Pretreatment with UbPA almost completely abolished subsequent labeling by FLAGUbPA, leaving only a faint residual band near 37 kDa, indicating effective inactivation of the UbPA-reactive enzyme population. In contrast, lysates pretreated with UbPA and then incubated with FLAGUbCMK retained prominent residual labeling. Notably, bands near approximately 25 kDa and a set of strongly intensified bands spanning the 50-100 kDa range were clearly observed. These residual bands correspond in molecular weight to the same species detected in untreated lysates labeled with FLAGUbCMK, shown in Figure 5A. This reflects that a substantial fraction of FLAGUbCMK-reactive proteins is not efficiently competed by UbPA pretreatment. This competition experiments demonstrate that FLAGUbCMK engages an additional subset of cysteine enzymes that are not susceptible to UbPA-mediated inactivation, underscoring the utility of CMK warhead in accessing enzyme populations that are refractory to alkyne-based activity profiling.

### Proteomic characterization of enzymes uniquely labeled by FLAGUbCMK

To characterize cysteine enzymes selectively labeled by FLAGUbCMK in UbPA-pretreated HEK293T cell lysates, protein-probe conjugates were enriched by immunoprecipitation with anti-FLAG magnetic agarose beads. The captured proteins were then subjected to on-bead trypsin digestion, and the resulting peptides were analyzed by liquid chromatography-tandem mass spectrometry (LC-MS/MS) for sequence assignment and protein identification. This proteomic analysis revealed robust and reproducible enrichment of a broad panel of E2 enzymes. In total, 22 E2 enzymes including UBE2D1, UBE2D2, UBE2D3, UBE2D4, UBE2E1, UBE2E2, UBE2E3, UBE2G1, UBE2G2, UBE2I, UBE2K, UBE2L3, UBE2L5, UBE2M, UBE2N, UBE2NL, UBE2O, UBE2S, UBE2T, UBE2V1, UBE2V2, and UFC1 were identified across multiple replicates. Of these, 17 E2s (UBE2D1, UBE2D2, UBE2D3, UBE2D4, UBE2E1, UBE2E2, UBE2E3, UBE2G1, UBE2K, UBE2L3, UBE2L5, UBE2M, UBE2N, UBE2NL, UBE2O, UBE2S, UBE2T) were robustly quantified across biological replicates and met the criteria for statistical testing and therefore appear as significantly enriched species in the volcano plots (Figure 5C). The remaining E2s (UBE2G2, UBE2I, UBE2V1, UBE2V2, and UFC1) were reproducibly identified but fall below the statistical threshold for P-value determination and are consequently not displayed in the volcano plots. All identified E2 enzymes are summarized in Table S3. Notably, these E2 enzymes remained highly enriched even after prior depletion by UbPA treatment, indicating that they are inefficiently captured by the classical UbPA probe and instead define an orthogonal population of E2 enzymes preferentially labeled by FLAGUbCMK. This selective enrichment underscores a mechanistic distinction between FLAGUbCMK and conventional DUB-targeting Ub probes, whereas UbPA is tuned to react with highly nucleophilic, low pKa active site cysteines characteristic to DUBs. E2 enzymes employ a higher pKa active site cysteine that are poorly engaged by an alkyne electrophile but are efficiently trapped by the more reactive CMK warhead. Apart from E2 enzymes for Ub, the presence of one UFM-specific E2 enzyme (UFC1) was also observed. Two E1 activating enzymes, UBA1 and UBA2, involved in Ub and SUMO pathways, respectively, and several DUBs including UCHL1, UCHL3, OTUB1, USP5, USP14, and OTUD6B were also identified. The presence and direct engagement of these DUBs were further validated by immunoblot analysis of immunoprecipitated FLAGUbCMK treated HEK293T lysates using DUB-specific antibodies (Figure S16). All identified proteins are listed in Table S3. A cross-interaction network and a depicting the relationship of these enzymes and their labeling by FLAGUbCMK probe are shown in Figure 5D.

Collectively, these results demonstrate that the FLAGUbCMK probe provides a powerful, chemoselective platform to map and quantify active population of E2 enzymes that are largely invisible to conventional Ub probes. The enrichment of multiple E2 enzymes following UbPA-mediated depletion establishes the complementary and non-overlapping reactivity profile of the CMK electrophile and underscores its utility as a chemical tool for broadening activity-based protein profiling coverage across the Ub signaling pathways.

### Validation of E2 Enzymes Identified from the Proteomic Analysis as FLAGUbCMK Targets

Our proteomic analysis revealed a discrete subset of E2 enzymes that were both strongly enriched and statistically significant relative to control samples. From this dataset, 10 E2s including UBE2D2, UBE2D3, UBE2E1, UBE2E3, UBE2K, UBE2L3, UBE2L5, UBE2M, UBE2N, and UBE2T emerged as high confidence targets. These hits span multiple E2 subfamilies, including canonical “workhorse” E2s (UBE2D2/3), the K48-linked chain-extending E2 UBE2K, a specialized K63 chain-building E2 (UBE2N), and E2s associated with distinct conjugation pathways (e.g., the NEDD8 E2 UBE2M and the Fanconi anemia E2 UBE2T), underscoring that FLAGUbCMK engages a mechanistically diverse but defined subset of E2 enzymes. To validate all six enzymes, we performed SDS-PAGE analysis of anti-FLAG immunoprecipitated HEK293T lysates pretreated with FLAGUbCMK, followed by western blotting with antibodies specific for UBE2D2, UBE2D3, UBE2E1, UBE2E3, UBE2K, UBE2L3, UBE2L5, UBE2M, UBE2N, and UBE2T. As shown in Figure 6A, clear enrichment of bands corresponding to covalent complexes between these enzymes and the FLAGUbCMK probe was observed. These bands were absent from control immunoprecipitates from untreated HEK293T lysates, underscoring the specificity and efficiency of the FLAGUbCMK probe in profiling distinct cysteine enzymes within the Ub signaling cascade.

**Figure 6:**
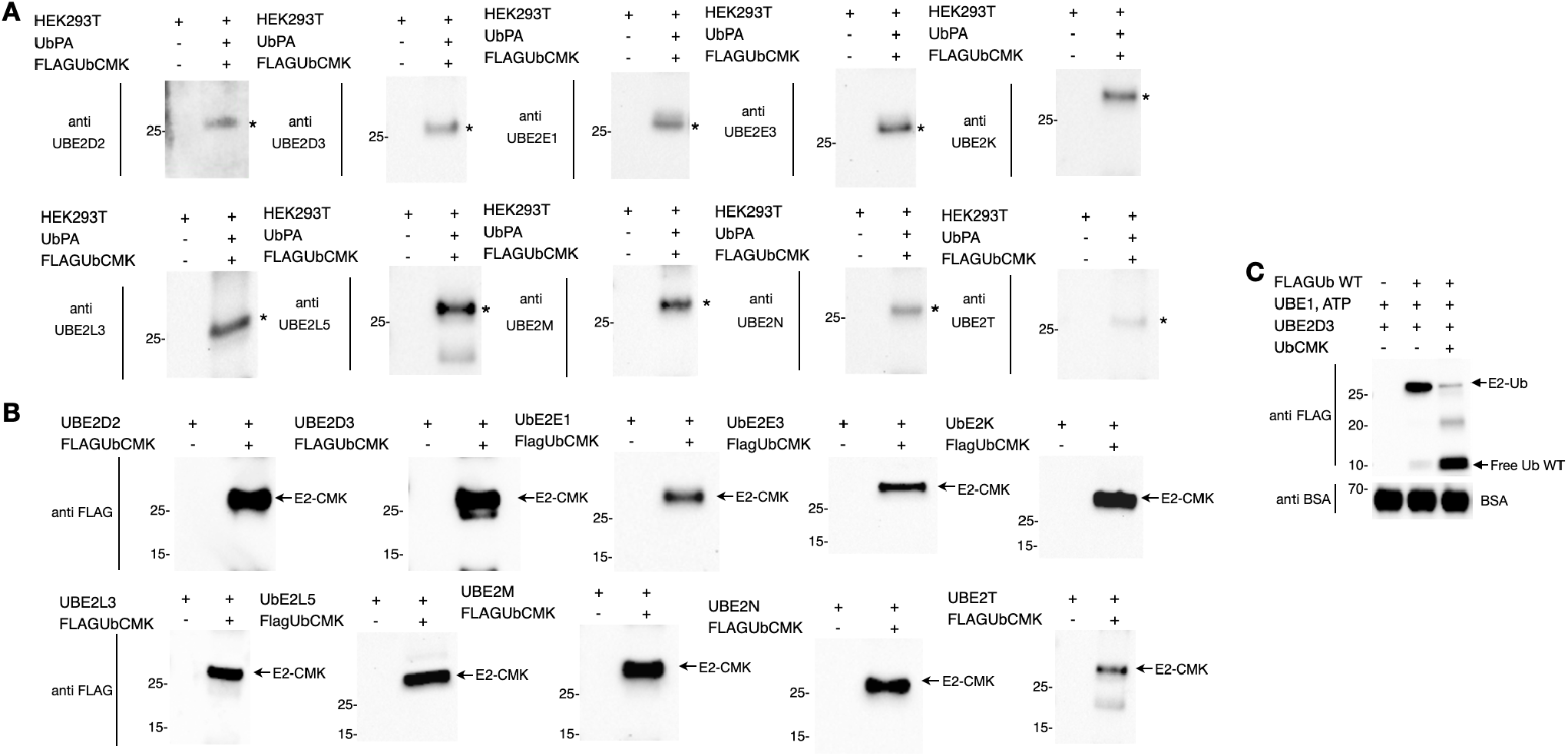
FLAGUbCMK probes covalently capture and functionally intercept E2 conjugating enzymes. (**A**) Validation of proteomics-identified E2 targets. SDS-PAGE and immunoblot analysis of the anti-FLAG immunoprecipitates using antibodies specific for UBE2D2, UBE2D3, UBE2E1, UBE2E3, UBE2K, UBE2L3, UBE2L5, UBE2M, UBE2N, and UBE2T. The asterisk indicates the enriched FLAGUbCMK-enzyme conjugates. (**B**) Recombinant UBE2D2, UBE2D3, UBE2E1, UBE2E3, UBE2K, UBE2L3, UBE2L5, UBE2M, UBE2N, and UBE2T were incubated with FLAGUbCMK *in vitro*. SDS-PAGE and immunoblotting with anti-FLAG antibody reveal discrete higher molecular weight E2-FLAGUbCMK adducts for all ten enzymes, with near-quantitative labeling for UBE2D2, UBE2D3, UBE2E1, UBE2E3, UBE2K, UBE2L3, UBE2L5, UBE2M, and UBE2N, and slower/partial labeling for UBE2T. (**C**) Inhibition of the E1-catalyzed E2-Ub formation by UbCMK. A reconstituted ubiquitination cascade containing UBE1, UBE2D3, ATP, and FLAG-ubiquitin (FLAGUb) was analyzed by anti-FLAG immunoblotting. In the absence of UbCMK, a prominent UBE2D3∼Ub band is observed, indicating efficient Ub transfer from E1 to E2. Addition of non-FLAG UbCMK suppresses E2∼Ub formation and leads to accumulation of free FLAGUb (∼10 kDa), consistent with covalent modification of the catalytic cysteine of the enzymes and blockade of thioester formation.

These 10 E2 enzymes were also independently validated as FLAGUbCMK-targeting enzymes in a biochemistry setup. We individually reacted each of the 10 E2 enzymes with FLAGUbCMK under defined *in vitro* conditions. Across this panel, all E2s formed covalent complexes with the probe, as evidenced by the appearance of discrete, higher molecular weight E2-FLAGUbCMK adducts by SDS-PAGE and immunoblotting (Figure 6B). The kinetics and extent of adduct formation varied among enzymes: UBE2D2, UBE2D3, UBE2E1, UBE2E3, UBE2K, UBE2L3, UBE2L5, UBE2M, and UBE2N all exhibited rapid and near-quantitative labeling under the conditions tested, whereas UBE2T showed slower or partial modification, consistent with differences in the active site microenvironment, intrinsic cysteine reactivity, and/or conformational accessibility of the catalytic residue.

The observation that all 10 proteomics-identified E2s are directly and covalently labeled by FLAGUbCMK confirms that their enrichment in complex lysates reflects *bona fide* probe engagement rather than indirect association within higher order protein complexes. At the same time, the enzyme-specific variation in labeling efficiency suggests that the CMK warhead is sufficiently versatile to accommodate structural diversity across E2 active sites, yet sensitive enough to report on underlying physicochemical differences within the E2 superfamily. Collectively, these findings establish that FLAGUbCMK can directly and covalently target a mechanistically relevant subset of E2 enzymes, thereby extending activity-based protein profiling of UPS enzymes beyond DUBs.

Prior work from Mulder *et al*. showed that UbDHA labels certain E2 enzymes via a catalysis-like acyl transfer mechanism.^11^ However, their systematic profiling also revealed that a subset of E2s, specifically UBE2F, UBE2I, UBE2L6, UBE2M, and UBE2Z, remained unreactive to UbDHA, even under conditions where other E2s were efficiently captured. This apparent “blind spot” suggested that certain E2 active sites are either less amenable to the UbDHA mechanism or present microenvironments that disfavor the conjugate-addition chemistry required for the dehydroalanine (DHA) engagement. UBE2M and UBE2I have already been discovered in our proteomic analysis of FLAGUbCMK-enriched proteins and UBEM2M reaction with FLAGUbCMK has been demonstrated in a biochemistry setup as shown in Figure S16. To confirm that other E2 enzymes that are nonreactive toward UbDHA can covalently bind to FLAGUbCMK, we tested their formation of covalent FLAGUbCMK complexes in a biochemistry setup as well. Recombinant UBE2F, UBE2I, UBE2L6, and UBE2Z were individually incubated with FLAGUbCMK under standard conditions, followed by SDS-PAGE and anti-FLAG immunoblotting to monitor covalent adduct formation. In contrast to their reported lack of reactivity toward UbDHA,^11^ all four E2s formed discrete higher molecular weight bands consistent with E2-FLAGUbCMK adducts (Figure S17). These results demonstrate that FLAGUbCMK can efficiently react with E2s and extend E2 coverage into a mechanistically important subset that was refractory to earlier probes.

### FLAGUbCMK as a Probe for E1 with an Active Site Cysteine

The proteomic analysis revealed both UBA1 and UBA2 that are E1 enzymes or components, indicating that FLAGUbCMK is an E1 probe as well and therefore UbCMK without a FLAG should function also as an E1 inhibitor. To test this, we reconstituted an E1-E2 ubiquitination cascade and conducted it with or without UbCMK. This reconstituted reaction cascade contained UBE1, ATP, UBE2D3, and FLAG-tagged Ub (FLAGUb). UbCMK was synthesized by ACPL from Ub_1_-_75_-Cys-6×His and confirmed by ESI-MS (Figure S18). Reactions were analyzed by immunoblotting with anti-FLAG to monitor formation of E2-Ub. In the absence of UbCMK, the cascade proceeded efficiently, yielding a robust FLAG-positive band at the apparent molecular weight of UBE2D3-Ub, consistent with successful transfer of Ub from UBE1 to UBE2D3 (Figure 6C). In contrast, inclusion of UbCMK in the reaction mixture markedly reduced formation of the UBE2D3-Ub complex. Under these conditions, anti-FLAG immunoblotting revealed pronounced accumulation of free FLAGUb at around 10 kDa, with concomitant loss of the E2-Ub band. In the canonical pathway, Ub is activated by E1 in an ATP-dependent reaction to form an E1-Ub thioester complex and then transferred to the active site cysteine of E2 by forming a E2-Ub thioester intermediate. We observed no UBE1-Ub formation, indicating inhibition of the E2-Ub formation was due to the inactivation of UBE1 by UbCMK. To test this prospect, we incubated UBE1 with FLAGUbCMK and analyzed the reaction by SDS–PAGE followed by anti-FLAG immunoblotting. We observed formation of a higher molecular weight FLAG-labeled species, consistent with covalent complex formation between UBE1 and FLAGUbCMK (Figure S19). These results strong support that FLAGUbCMK is an E1 activity probe as well.

### FLAGUbCMK as a Probe to Quantify Active E2 Enzymes in Different Stress Conditions

To evaluate the utility of FLAGUbCMK for profiling E2 enzymes under biologically relevant conditions, we examined its ability to capture endogenous E2s from HEK293T cells subjected to distinct forms of cellular stress. We focused on oxidative, proteotoxic, inflammatory, and metabolic stress, as well as lipid oxidation and genotoxic stress, each of which is known to selectively mobilize specific arms of the UPS. In all cases, whole cell lysates were incubated with FLAGUbCMK, followed by anti-FLAG immunoprecipitation and Western blotting with different anti-E2 antibodies. BSA was used as an exogenous loading control. We reasoned that if FLAGUbCMK faithfully captures catalytically engaged E2s, stress-induced E2 mobilization should be reflected as increased probe-dependent pull-down relative to unstressed controls.

Under oxidative stress, induced by treating HEK293T cells with the thiol-oxidizing agent diamide (400 µM, 20 min), we observed a marked increase in global protein ubiquitination, consistent with activation of Ub-dependent protein quality-control pathways. We monitored UBE2M, a stress-inducible NEDD8-conjugating E2 that functions dually as a cullin neddylation enzyme under basal conditions and as a Ub E2 in the Parkin-DJ-1 stress-response axis.^26^ As shown in Figure 7A, Anti-UBE2M immunoblotting of FLAGUbCMK pulldowns revealed substantially enhanced capture of Ub-UBE2M conjugates from diamide-treated lysates compared to untreated controls, indicating oxidative stress dependent mobilization of catalytically engaged UBE2M.

**Figure 7:**
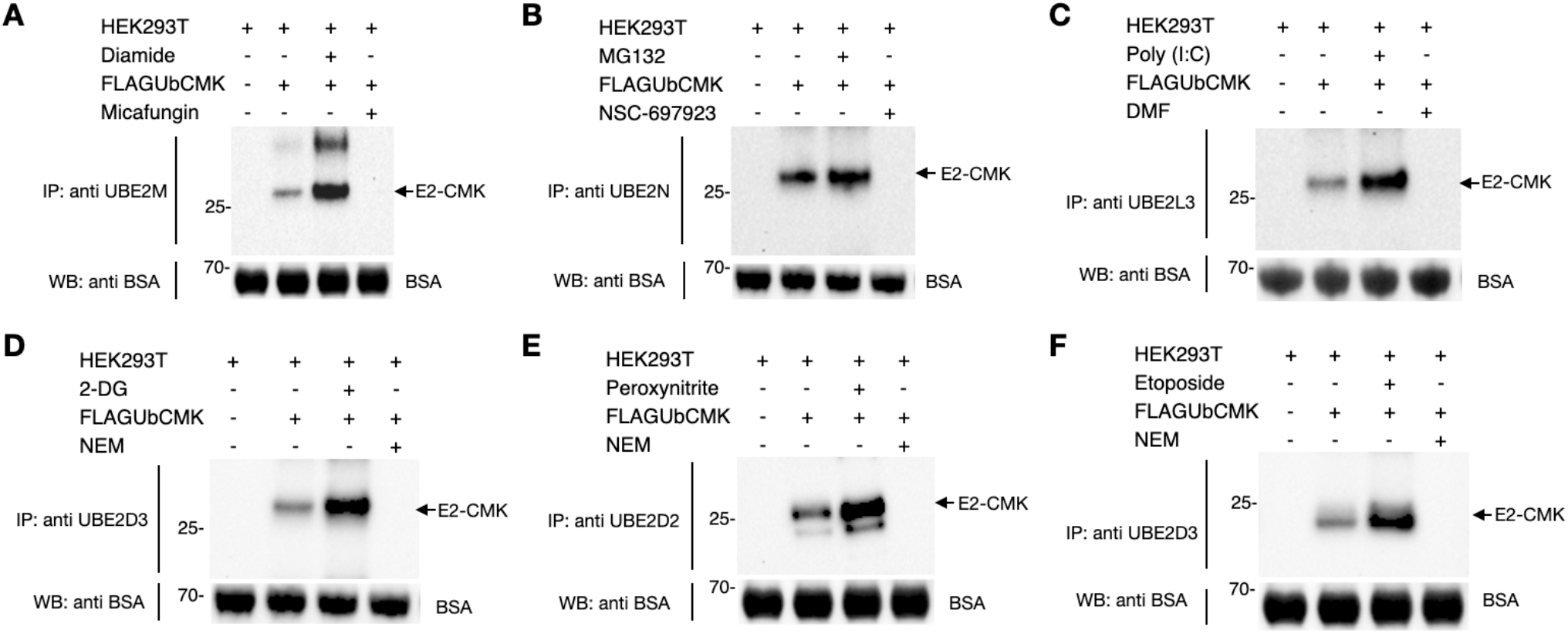
Stress-dependent activity-based capture of endogenous E2 conjugating enzymes by FLAGUbCMK. HEK293T cells were subjected to (**A**) oxidative stress (400 µM diamide, 20 min), (**B**) proteotoxic stress (MG132), (**C**) inflammatory signaling [poly(I:C)], (**D**) metabolic stress (10 mM 2-deoxy-D-glucose, 2-DG), (**E**) lipid oxidative stress (oxidized LDL generated by peroxynitrite), or (**F**) genotoxic stress by etoposide (DNA-damaging agent), followed by lysis and incubation with FLAGUbCMK. Probe-labeled proteins were enriched by anti-FLAG immunoprecipitation and analyzed by immunoblotting with E2-specific antibodies against UBE2M (oxidative), UBE2N (proteotoxic), UBE2L3 (inflammatory), UBE2D3 (metabolic and DNA damage), and UBE2D2 (lipid oxidative stress). BSA was used as a loading control. Stress treatments enhanced FLAGUbCMK capture of the corresponding E2s relative to untreated controls, indicating increased catalytic engagement under each stress condition. For UBE2M, UBE2N, and UBE2L3, pre-treatment with Micafungin, NSC-697923, or dimethyl fumarate (DMF), respectively, abolished probe labeling, confirming active-site dependent capture. For UBE2D2 and UBE2D3, pre-alkylation with *N*-ethylmaleimide (NEM) similarly blocked FLAGUbCMK labeling, demonstrating dependence on accessible catalytic cysteines.

Proteotoxic stress was modeled using the proteasome inhibitor MG132, which blocks proteasomal degradation, leading to accumulation of polyubiquitinated substrates and activation of compensatory Ub-dependent quality-control circuits.^27^ Consistent with this, MG132 treatment increased overall polyUb levels. We monitored UBE2N (Ubc13), the sole E2 that exclusively catalyzes K63-linked polyubiquitin chain formation, which plays a central non-degradative signaling role in protein quality control.^28^ As observed in Figure 7B, FLAGUbCMK pulldowns from MG132-treated lysates exhibited substantially higher Ub-UBE2N capture relative to untreated controls, confirming that the probe reports on proteotoxic stress induced engagement of the UBE2N.

Inflammatory signaling was induced by polyinosinic:polycytidylic acid [poly(I:C)], a synthetic double-stranded RNA analog that activates TLR3-dependent NF-κB signaling and proinflammatory cytokine production.^29^ In this context, we monitored UBE2L3 (UbcH7), a preferred E2 for LUBAC-mediated NF-κB activation.^30^ Poly(I:C) treatment led to a pronounced increase in FLAGUbCMK-dependent UBE2L3 capture (Figure 7C), indicating that inflammatory signaling enhances the pool of catalytically accessible UBE2L3 that can be covalently trapped by FLAGUbCMK.

We next investigated E2 mobilization in a metabolic stress setting. Tumor cells such as glioma frequently operate under chronic nutrient and energy imbalance driven by high glycolytic demand, and UBE2D3 has been implicated in this context, where it promotes SHP-2 ubiquitination and STAT3-driven glycolysis.^31^ To model tumor-like metabolic stress in HEK293T cells, we treated cells with 2-deoxy-D-glucose (2-DG, 10 mM), a glycolytic inhibitor that disrupts ATP production and induces a glycolysis-linked stress response. In this system, 2-DG treatment impaired glycolytic ATP generation and activated adaptive stress pathways reminiscent of those seen in highly glycolytic glioma cells. Monitoring UBE2D3 as a readout of metabolic stress associated E2 mobilization, we observed enhanced Ub-UBE2D3 capture by FLAGUbCMK from 2-DG-treated lysates relative to untreated cells (Figure 7D), demonstrating that the probe detects metabolic-stress-induced engagement of this E2.

We further extended this analysis to additional stress paradigms relevant to UBE2D family enzymes. LDL oxidation by peroxynitrite was used to generate oxidized LDL (oxLDL), a model of lipid oxidative stress known to perturb redox balance and trigger adaptive Ub signaling.^32^ Under oxLDL conditions, we monitored UBE2D2 and observed increased capture of Ub-UBE2D2 adducts by FLAGUbCMK (Figure 7E), indicating that UBE2D2 is mobilized in response to oxidative lipid stress and is accessible to covalent trapping by the probe. In parallel, we examined UBE2D3 mobilization upon DNA damage induction in HEK293T cells, using a genotoxic stressor (etoposide) known to activate DNA damage response pathways. UBE2D3 facilitates DNA damage response through nonhomologous end joining.^33^ Under these DNA damage conditions, as shown in Figure 7F, FLAGUbCMK pulldowns revealed elevated Ub-UBE2D3 capture relative to untreated controls, consistent with stress dependent engagement of UBE2D3 in genotoxic signaling.

Because specific small molecule inhibitors are not available for UBE2D2 and UBE2D3, we employed a global thiol blocking strategy to validate that probe labeling of these E2s reflects active site modification. Pretreatment of HEK293T cells with *N*-ethylmaleimide (NEM), a broad cysteine alkylator, before FLAGUbCMK incubation completely abolished FLAGUbCMK-dependent capture of UBE2D2 and UBE2D3. This loss of labeling demonstrates that probe engagement requires accessible cysteine residues, consistent with the expected modification of the catalytic cysteine. For UBE2M, UBE2N, and UBE2L3, we carried out inhibitor-based competition experiments to further confirm that FLAGUbCMK labeling reflects catalytic engagement rather than nonspecific binding. Micafungin, an echinocandin antifungal recently shown to inhibit UBE2M-dependent neddylation, was used to block UBE2M activity.^34^ NSC-697923, a covalent inhibitor that alkylates the catalytic cysteine of UBE2N and abolishes K63-linked ubiquitination, was used to inhibit UBE2N.^35^ And dimethyl fumarate (DMF), which modifies catalytic cysteines in E2 enzymes and suppresses UBE2L3-mediated inflammatory signaling, was used to target UBE2L3.^36^ In each case, pretreatment with the corresponding inhibitor completely abolished FLAGUbCMK capture of the targeted E2 (Figure 7A-F), consistent with the requirement for an unmodified, catalytically competent active-site cysteine.

Taken together, our data show that FLAGUbCMK selectively and sensitively reports on the mobilization of E2 enzymes across multiple mechanistically distinct stress conditions, establishing FLAGUbCMK as a robust activity-based probe for E2 enzymes and potentiating stress-responsive, activity-dependent profiling of cysteine enzymes in the UPS under diverse physiological and pathophysiological conditions.

## CONCLUSION

E2s occupy a central catalytic position in Ub signaling, however, tools for directly probing their activity have remained less developed than those available for DUBs. This limitation stems from the relatively low nucleophilicity of E2 active site cysteines, which makes them difficult to covalently capture using conventional Ub probes. In this work, w addressed this long-standing gap by developing FLAGUbCMK as a standalone, E1-independent activity-based probe that irreversibly engages E2 catalytic cysteines. Guided by DFT calculations, we identify CMK with a low activation barrier for reaction with weakly nucleophilic thiolates, providing a mechanistic rationale for its compatibility for E2 bioconjugation. FLAGUbCMK was synthesis via ACPL and underwent irreversible covalent engagement of E2s, E1s, and DUBs. Its use as an active-based probe in HEK293T cell lysates captured multiple E2 enzyme subfamilies with broad coverage. It also supports activity-dependent quantification of endogenous E2 mobilization under diverse physiological stress conditions. These capabilities establish FLAGUbCMK as a versatile chemical probe for monitoring dynamic changes in Ub transfer capacity under biologically relevant perturbations.

Since FLAGUbCMK also engages other cysteine-dependent components in the UPS, including E1s and DUBs, this broader reactivity positions FLAGUbCMK as a versatile chemical handle for capturing multiple catalytic nodes within Ub signaling pathway. Overall, FLAGUbCMK provides a simple and powerful solution for covalent interrogation of E2 enzymes and expands activity-based profiling coverage across the UPS, enabling both mechanistic and functional studies of E2 biology in both physiological and disease-relevant contexts.

## ASSOCIATED CONTENT

### Notes

The authors declare no competing financial interests. All data supporting the findings in this article are available in the main text or the supplementary information.

## SUPPORTING INFORMATION

Experimental procedures for protein expression, Ub probe synthesis, characterization of probes and their reactions with different enzymes, protein profiling using synthesized probes from cell lysates, validation of identified enzymes from proteomic analysis, stress dependent activity profiling, supplementary figures and tables.

## ACKNOWLEDGMENT

This work was partially supported by the Welch Foundation (grant A-1715 to W.R.L.), National Institutes of Health (grants R35GM145351 and R01CA291968 to W.R.L.), Cancer Prevention and Research Institute of Texas (grant RP230345 to W.R.L.), and the Texas A&M Advancing Discovery to Market Program. The authors thank Dr. Yohannes Rezenom in the Mass Spectrometry Facility of Texas A&M University for helping with running the LC-MS characterizations of all proteins.

## Insert Table of Contents artwork here

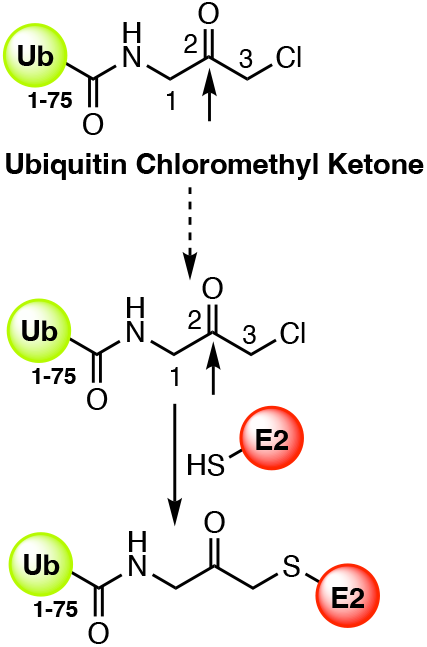

## REFERNCES

(1) Swatek, K. N.; Komander, D. Ubiquitin modifications. Cell Research 2016, 26 (4), 399–422.

(2) Komander, D.; Rape, M. The ubiquitin code. Annu Rev Biochem 2012, 81, 203–229.

(3) Trempe, J. F. Reading the ubiquitin postal code. Curr Opin Struct Biol 2011, 21 (6), 792–801.

(4) Ciechanover, A. The ubiquitin-proteasome pathway: on protein death and cell life. EMBO J. 1998, 17 (24), 7151–7160.

(5) Ciechanover, A.; Schwartz, A. L. The ubiquitin-proteasome pathway: the complexity and myriad functions of proteins death. Proc. Natl. Acad. Sci. U. S. A. 1998, 95 (6), 2727–2730.

(6) Schwartz, A. L.; Ciechanover, A. The ubiquitin-proteasome pathway and pathogenesis of human diseases. Annu. Rev. Med. 1999, 50, 57–74. Hegde, A. N.; Upadhya, S. C. The ubiquitin-proteasome pathway in health and disease of the nervous system. Trends Neurosci. 2007, 30 (11), 587–595. Wang, J.; Maldonado, M. A. The ubiquitin-proteasome system and its role in inflammatory and autoimmune diseases. Cell. Mol. Immunol. 2006, 3 (4), 255–261. Meyer-Schwesinger, C. The ubiquitin-proteasome system in kidney physiology and disease. Nat Rev Nephrol 2019, 15 (7), 393–411. Oddo, S. The ubiquitin-proteasome system in Alzheimer’s disease. J Cell Mol Med 2008, 12 (2), 363–373.

(7) Knoll, N.; Neamati, N.; Üren, A. Mechanisms and Design Principles of Proteolysis-Targeting Chimeras and Their Emerging Applications. ACS Pharmacol Transl Sci 2026, 9 (4), 784–814. Békés, M.; Langley, D. R.; Crews, C. M. PROTAC targeted protein degraders: the past is prologue. Nature Reviews Drug Discovery 2022, 21 (3), 181–200.

(8) Wang, T.; Yin, L.; Cooper, E. M.; Lai, M. Y.; Dickey, S.; Pickart, C. M.; Fushman, D.; Wilkinson, K. D.; Cohen, R. E.; Wolberger, C. Evidence for bidentate substrate binding as the basis for the K48 linkage specificity of otubain 1. J Mol Biol 2009, 386 (4), 1011–1023. Hershko, A.; Rose, I. A. Ubiquitin-aldehyde: a general inhibitor of ubiquitin-recycling processes. Proceedings of the National Academy of Sciences 1987, 84 (7), 1829–1833.

(9) Lam, Y. A.; Xu, W.; DeMartino, G. N.; Cohen, R. E. Editing of ubiquitin conjugates by an isopeptidase in the 26S proteasome. Nature 1997, 385 (6618), 737–740.

(10) Hemelaar, J.; Galardy, P. J.; Borodovsky, A.; Kessler, B. M.; Ploegh, H. L.; Ovaa, H. Chemistry-based functional proteomics: mechanism-based activity-profiling tools for ubiquitin and ubiquitin-like specific proteases. J. Proteome Res. 2004, 3 (2), 268–276.

(11) Mulder, M. P.; Witting, K.; Berlin, I.; Pruneda, J. N.; Wu, K. P.; Chang, J. G.; Merkx, R.; Bialas, J.; Groettrup, M.; Vertegaal, A. C.; et al. A cascading activity-based probe sequentially targets E1-E2-E3 ubiquitin enzymes. Nat. Chem. Biol. 2016, 12 (7), 523–530.

(12) Ekkebus, R.; van Kasteren, S. I.; Kulathu, Y.; Scholten, A.; Berlin, I.; Geurink, P. P.; de Jong, A.; Goerdayal, S.; Neefjes, J.; Heck, A. J. R.; et al. On Terminal Alkynes That Can React with Active-Site Cysteine Nucleophiles in Proteases. Journal of the American Chemical Society 2013, 135 (8), 2867–2870. Love, K. R.; Pandya, R. K.; Spooner, E.; Ploegh, H. L. Ubiquitin C-terminal electrophiles are activity-based probes for identification and mechanistic study of ubiquitin conjugating machinery. ACS Chem. Biol. 2009, 4 (4), 275–287. Pao, K. C.; Stanley, M.; Han, C.; Lai, Y. C.; Murphy, P.; Balk, K.; Wood, N. T.; Corti, O.; Corvol, J. C.; Muqit, M. M.; et al. Probes of ubiquitin E3 ligases enable systematic dissection of parkin activation. Nat. Chem. Biol. 2016, 12 (5), 324–331. Gong, L.; Kamitani, T.; Millas, S.; Yeh, E. T. H. Identification of a Novel Isopeptidase with Dual Specificity for Ubiquitin- and NEDD8-conjugated Proteins *. Journal of Biological Chemistry 2000, 275 (19), 14212–14216. Gong, P.; Davidson, G. A.; Gui, W.; Yang, K.; Bozza, W. P.; Zhuang, Z. Activity-based ubiquitin-protein probes reveal target protein specificity of deubiquitinating enzymes. Chemical Science 2018, 9 (40), 7859–7865, 10.1039/C8SC01573B. Gui, W.; Ott, C. A.; Yang, K.; Chung, J. S.; Shen, S.; Zhuang, Z. Cell-Permeable Activity-Based Ubiquitin Probes Enable Intracellular Profiling of Human Deubiquitinases. J. Am. Chem. Soc. 2018, 140 (39), 12424–12433. Meledin, R.; Mali, S. M.; Kleifeld, O.; Brik, A. Activity-Based Probes Developed by Applying a Sequential Dehydroalanine Formation Strategy to Expressed Proteins Reveal a Potential α-Globin-Modulating Deubiquitinase. i>Angew Chem Int Ed Engl 2018, 57 (20), 5645–5649. Xu, L.; Fan, J.; Wang, Y.; Zhang, Z.; Fu, Y.; Li, Y.-M.; Shi, J. An activity-based probe developed by a sequential dehydroalanine formation strategy targets HECT E3 ubiquitin ligases. Chemical Communications 2019, 55 (49), 7109–7112, 10.1039/C9CC03739J. Gui, W.; Shen, S.; Zhuang, Z. Photocaged Cell-Permeable Ubiquitin Probe for Temporal Profiling of Deubiquitinating Enzymes. J. Am. Chem. Soc. 2020, 142 (46), 19493–19501. Liang, L.-J.; Chu, G.-C.; Qu, Q.; Zuo, C.; Mao, J.; Zheng, Q.; Chen, J.; Meng, X.; Jing, Y.; Deng, H.; et al. Chemical Synthesis of Activity-Based E2-Ubiquitin Probes for the Structural Analysis of E3 Ligase-Catalyzed Transthiolation. Angewandte Chemie International Edition 2021, 60 (31), 17171–17177. Wang, Y.; Chen, J.; Hua, X.; Meng, X.; Cai, H.; Wang, R.; Shi, J.; Deng, H.; Liu, L.; Li, Y. M. Photocaging of Activity-Based Ubiquitin Probes via a C-Terminal Backbone Modification Strategy. Angew. Chem. Int. Ed. 2022, 61 (28), e202203792.

(13) Mevissen, T. E.; Hospenthal, M. K.; Geurink, P. P.; Elliott, P. R.; Akutsu, M.; Arnaudo, N.; Ekkebus, R.; Kulathu, Y.; Wauer, T.; El Oualid, F.; et al. OTU deubiquitinases reveal mechanisms of linkage specificity and enable ubiquitin chain restriction analysis. Cell 2013, 154 (1), 169–184, Research Support, Non-U.S. Gov’t.

(14) Chanda, S.; Atla, S.; Sheng, X.; Nyalata, S.; Alugubelli, Y. R.; Coleman, D. D.; Jiang, W.; Lopes, R.; Guo, S.; Wand, A. J.; et al. Ubiquitin Azapeptide Esters as Next-Generation Activity-Based Probes for Cysteine Enzymes in the Ubiquitin Signal Pathway. Journal of the American Chemical Society 2025, 147 (21), 17817–17828. Chanda, S.; Liu, W. R. From Covalent Traps to Fluorescent Beacons: The Expanding Arsenal of Chemical Probes for Studying Ubiquitin and Ubiquitin-Like Proteins. Angew Chem Int Ed Engl 2026, e20118.

(15) Chanda, S.; Pham, A.; Karnati, S.; Rodriguez, N. S.; Toner, A.; Liu, W. R. Synthetic Strategies for Activity-Based Probes to Decode Ubiquitin-Like Modifiers. Chemistry (Easton) 2026, e03597.

(16) Parashar, S.; Kaushik, A.; Ambasta, R. K.; Kumar, P. E2 conjugating enzymes: A silent but crucial player in ubiquitin biology. Ageing Res Rev 2025, 108, 102740. Stewart, M. D.; Ritterhoff, T.; Klevit, R. E.; Brzovic, P. S. E2 enzymes: more than just middle men. Cell Res. 2016, 26 (4), 423–440.

(17) Guo, Y.; Chen, X.; Zhang, X.; Hu, X. UBE2S and UBE2C confer a poor prognosis to breast cancer via downregulation of Numb. Front. Oncol. 2023, 13, 992233.

(18) Machida, Y. J.; Machida, Y.; Chen, Y.; Gurtan, A. M.; Kupfer, G. M.; D’Andrea, A. D.; Dutta, A. UBE2T is the E2 in the Fanconi anemia pathway and undergoes negative autoregulation. Mol. Cell 2006, 23 (4), 589–596. Watanabe, A.; Lu, J.; Ishihara, K.; Iwabuchi, S.; Ohno, K.; Hashimoto, S.; Asakage, T.; Takahashi, K.; Podyma-Inoue, K. A.; Watabe, T. UBE2T promotes epithelial-mesenchymal transition and motility in oral cancer cells via induction of IL-6 expression. Oncol. Lett. 2025, 30 (4), 473. Yu, Z.; Jiang, X.; Qin, L.; Deng, H.; Wang, J.; Ren, W.; Li, H.; Zhao, L.; Liu, H.; Yan, H.; et al. A novel UBE2T inhibitor suppresses Wnt/beta-catenin signaling hyperactivation and gastric cancer progression by blocking RACK1 ubiquitination. Oncogene 2021, 40 (5), 1027–1042. Ma, N.; Li, Z.; Yan, J.; Liu, X.; He, L.; Xie, R.; Lu, X. Diverse roles of UBE2T in cancer (Review). Oncol. Rep. 2023, 49 (4).

(19) Lenoir, J. J.; Parisien, J. P.; Horvath, C. M. Immune regulator LGP2 targets Ubc13/UBE2N to mediate widespread interference with K63 polyubiquitination and NF-kappaB activation. Cell Rep. 2021, 37 (13), 110175. Ishikawa, C.; Barreyro, L.; Sampson, A. M.; Hueneman, K. M.; Choi, K.; Philbrook, S. Y.; Choi, I.; Bolanos, L. C.; Wunderlich, M.; Volk, A. G.; et al. Ubiquitin-conjugating enzyme UBE2N modulates proteostasis in immunoproteasome-positive acute myeloid leukemia. J. Clin. Invest. 2025, 135 (10).

(20) Tolbert, B. S.; Tajc, S. G.; Webb, H.; Snyder, J.; Nielsen, J. E.; Miller, B. L.; Basavappa, R. The active site cysteine of ubiquitin-conjugating enzymes has a significantly elevated pKa: functional implications. Biochemistry 2005, 44 (50), 16385–16391.

(21) Delos Reyes, A. M. V.; Lux, M. C.; Hann, Z. S.; Ji, C.; Kochanczyk, T.; DiBello, M.; Lima, C. D.; Tan, D. S. Design and Semisynthesis of Biselectrophile-Functionalized Ubiquitin Probes To Investigate Transthioesterification Reactions. Org. Lett. 2024, 26 (22), 4594–4599.

(22) Pruneda, J. N.; Stoll, K. E.; Bolton, L. J.; Brzovic, P. S.; Klevit, R. E. Ubiquitin in motion: structural studies of the ubiquitin-conjugating enzyme∼ubiquitin conjugate. Biochemistry 2011, 50 (10), 1624–1633. Pickart, C. M.; Eddins, M. J. Ubiquitin: structures, functions, mechanisms. Biochimica et Biophysica Acta (BBA) - Molecular Cell Research 2004, 1695 (1), 55–72.

(23) Otto, H. H.; Schirmeister, T. Cysteine Proteases and Their Inhibitors. Chem. Rev. 1997, 97 (1), 133–172. Malthouse, J. P. G.; Mackenzie, N. E.; Boyd, A. S. F.; Scott, A. I. Detection of a Tetrahedral Adduct in a Trypsin-Chloromethyl Ketone Specific Inhibitor Complex by C-13 Nmr. Journal of the American Chemical Society 1983, 105 (6), 1685–1686.

(24) Muir, T. W.; Sondhi, D.; Cole, P. A. Expressed protein ligation: A general method for protein engineering. Proceedings of the National Academy of Sciences 1998, 95 (12), 6705–6710. Zhao, B.; Tsai, Y. C.; Jin, B.; Wang, B.; Wang, Y.; Zhou, H.; Carpenter, T.; Weissman, A. M.; Yin, J. Protein Engineering in the Ubiquitin System: Tools for Discovery and Beyond. Pharmacol. Rev. 2020, 72 (2), 380–413.

(25) Chanda, S.; Pham, A.; Shi, Y.; Atla, S.; Liu, W. R. A Simple, Quick, and Scalable Route to Fluorogenic Ubiquitin and Ubiquitin-Like Protein Substrates for Assessing Activities of Deubiquitinases and Ubiquitin-Like Protein-Specific Proteases. ACS Chemical Biology 2025. Qiao, Y.; Yu, G.; Kratch, K. C.; Wang, X. A.; Wang, W. W.; Leeuwon, S. Z.; Xu, S.; Morse, J. S.; Liu, W. R. Expressed Protein Ligation without Intein. J. Am. Chem. Soc. 2020, 142 (15), 7047–7054. Yu, G.; Qiao, Y.; Blankenship, L. R.; Liu, W. R. Protein Synthesis via Activated Cysteine-Directed Protein Ligation. Methods Mol. Biol. 2022, 2530, 159–167.

(26) Zhou, W.; Xu, J.; Tan, M.; Li, H.; Li, H.; Wei, W.; Sun, Y. UBE2M Is a Stress-Inducible Dual E2 for Neddylation and Ubiquitylation that Promotes Targeted Degradation of UBE2F. Mol Cell 2018, 70 (6), 1008–1024.e1006. Zhou, L.; Lin, X.; Zhu, J.; Zhang, L.; Chen, S.; Yang, H.; Jia, L.; Chen, B. NEDD8-conjugating enzyme E2s: critical targets for cancer therapy. Cell Death Discovery 2023, 9 (1), 23. Xiong, H.; Wang, D.; Chen, L.; Choo, Y. S.; Ma, H.; Tang, C.; Xia, K.; Jiang, W.; Ronai, Z.; Zhuang, X.; et al. Parkin, PINK1, and DJ-1 form a ubiquitin E3 ligase complex promoting unfolded protein degradation. J Clin Invest 2009, 119 (3), 650–660.

(27) Pai, J. T.; Chen, L. P.; Chang, H. J.; Wang, S. W.; Leu, Y. L.; Lai, C. T.; Weng, M. S. Proteostasis Disruption by Proteasome Inhibitor MG132 and Propolin G Induces ER Stress- and Autophagy-Mediated Apoptosis in Breast Cancer. Food Sci Nutr 2025, 13 (7), e70632.

(28) Hodge, C. D.; Spyracopoulos, L.; Glover, J. N. Ubc13: the Lys63 ubiquitin chain building machine. Oncotarget 2016, 7 (39), 64471–64504. Eddins, M. J.; Carlile, C. M.; Gomez, K. M.; Pickart, C. M.; Wolberger, C. Mms2–Ubc13 covalently bound to ubiquitin reveals the structural basis of linkage-specific polyubiquitin chain formation. Nature Structural & Molecular Biology 2006, 13 (10), 915–920.

(29) Stowell, N. C.; Seideman, J.; Raymond, H. A.; Smalley, K. A.; Lamb, R. J.; Egenolf, D. D.; Bugelski, P. J.; Murray, L. A.; Marsters, P. A.; Bunting, R. A.; et al. Long-term activation of TLR3 by poly(I:C) induces inflammation and impairs lung function in mice. Respir Res 2009, 10 (1), 43.

(30) Lewis, M. J.; Vyse, S.; Shields, A. M.; Boeltz, S.; Gordon, P. A.; Spector, T. D.; Lehner, P. J.; Walczak, H.; Vyse, T. J. UBE2L3 polymorphism amplifies NF-κB activation and promotes plasma cell development, linking linear ubiquitination to multiple autoimmune diseases. Am J Hum Genet 2015, 96 (2), 221–234.

(31) Pan, Z.; Bao, J.; Zhang, L.; Wei, S. UBE2D3 Activates SHP-2 Ubiquitination to Promote Glycolysis and Proliferation of Glioma via Regulating STAT3 Signaling Pathway. Front Oncol 2021, 11, 674286.

(32) Lara-Guzmán, O. J.; Gil-Izquierdo, Á.; Medina, S.; Osorio, E.; Álvarez-Quintero, R.; Zuluaga, N.; Oger, C.; Galano, J. M.; Durand, T.; Muñoz-Durango, K. Oxidized LDL triggers changes in oxidative stress and inflammatory biomarkers in human macrophages. Redox Biol 2018, 15, 1–11. Roman-Trufero, M.; Dillon, N. The UBE2D ubiquitin conjugating enzymes: Potential regulatory hubs in development, disease and evolution. Frontiers in Cell and Developmental Biology 2022, Volume 10 - 2022, Review.

(33) Yalçin, Z.; Lam, S. Y.; Peuscher, M. H.; van der Torre, J.; Zhu, S.; Iyengar, P. V.; Salas-Lloret, D.; de Krijger, I.; Moatti, N.; van der Lugt, R.; et al. UBE2D3 facilitates NHEJ by orchestrating ATM signalling through multi-level control of RNF168. Nat Commun 2024, 15 (1), 5032. Yang, H.; Wu, L.; Ke, S.; Wang, W.; Yang, L.; Gao, X.; Fang, H.; Yu, H.; Zhong, Y.; Xie, C.; et al. Downregulation of Ubiquitin-conjugating Enzyme UBE2D3 Promotes Telomere Maintenance and Radioresistance of Eca-109 Human Esophageal Carcinoma Cells. J Cancer 2016, 7 (9), 1152–1162. Yalçin, Z.; Koot, D.; Bezstarosti, K.; Salas-Lloret, D.; Bleijerveld, O. B.; Boersma, V.; Falcone, M.; González-Prieto, R.; Altelaar, M.; Demmers, J. A. A.; et al. Ubiquitinome Profiling Reveals in Vivo UBE2D3 Targets and Implicates UBE2D3 in Protein Quality Control. Mol Cell Proteomics 2023, 22 (6), 100548.

(34) Mamun, M. A. A.; Liu, S.; Zhao, L.; Zhao, L.; Li, Z. R.; Shen, D.; Zheng, Y.; Zheng, Y. C.; Liu, H. M. Micafungin: A promising inhibitor of UBE2M in cancer cell growth suppression. Eur. J. Med. Chem. 2023, 260, 115732.

(35) Cheng, J.; Fan, Y. H.; Xu, X.; Zhang, H.; Dou, J.; Tang, Y.; Zhong, X.; Rojas, Y.; Yu, Y.; Zhao, Y.; et al. A small-molecule inhibitor of UBE2N induces neuroblastoma cell death via activation of p53 and JNK pathways. Cell Death Dis 2014, 5 (2), e1079.

(36) McGuire, V. A.; Ruiz-Zorrilla Diez, T.; Emmerich, C. H.; Strickson, S.; Ritorto, M. S.; Sutavani, R. V.; Weiβ, A.; Houslay, K. F.; Knebel, A.; Meakin, P. J.; et al. Dimethyl fumarate blocks pro-inflammatory cytokine production via inhibition of TLR induced M1 and K63 ubiquitin chain formation. Scientific Reports 2016, 6 (1), 31159.

